# Gene specificity landscapes for comparative transcriptomic analysis across tissues, cell types, and species

**DOI:** 10.1101/2025.06.04.657813

**Authors:** Erik Bot, Jose Davila-Velderrain

## Abstract

Gene expression specificity is a biological parameter relevant for understanding the molecular basis of evolutionary constraints and tissue-selective pathogenesis. Many efforts have tried to quantify the degree of tissue specificity of individual genes. The growing availability of single-cell transcriptomic data greatly expands the context in which expression specificity can be assessed. We present a computational strategy to globally analyse and compare the specificity of genes and groups of related genes across different contexts. By representing expression profiles in terms of expression level-breadth (L-B) relationships, we are able to quantify and construct 2D landscapes that provide a globally consistent coordinate system to map specificity patterns. We characterize these landscapes at different levels of resolution and across species to demonstrate simple strategies for comparative transcriptomics. We use this approach to investigate the tissue, cell type, and neuronal specificity of human genes and generate reference specificity landscapes. Finally, by comparing the specificity of brain cell subtypes across 4 primate species we find that their degree of conservation mirrors evolutionary divergence times. Our analysis framework and data resources are available in the R package *GeneSLand*.

## Introduction

A cell’s function is mediated at the molecular level by the kind and number of molecules it expresses. Gene regulation allows cells to differentiate and express different genes despite a virtually invariant genome^1^. By selectively switching on and off genes, cells can diversify their molecular machineries and perform specialized functions while maintaining core processes. Profiling what genes are on or off in a given condition thus offers an operational means to define functionally relevant cell states, and the degree to which genes are on or off across conditions reveals valuable information about gene function and regulatory principles^2,3^. The introduction of genome-wide transcriptomic analysis provided a systematic way to estimate cell states and analyse expression patterns^4–6^. Whether a gene is expressed or not in a given context is an intuitive question from biologists. However, empirically defining on-off gene activity configurations from high-throughput experiments turned out to be not so straightforward. Different genes are expressed at different levels in any given context, and these levels commonly vary for the same genes across contexts. This, together with signal-to-noise variability among measurement technologies and heterogeneity of source tissue, contributes to the infeasibility of simply discretizing gene expression levels to determine expression status^7^. Single-cell transcriptomic technologies seem at first sight to bypass some of these problems by directly estimating what is expressed or not in each cell. In practice, however, the inherent sparsity of the data generated with these technologies does not allow a straightforward distinction between biological or technical zero values^8,9^. Despite these problems, expression profiles have been extensively used to investigate patterns of cell and tissue expression either by defining absolute threshold values within individual samples or by estimating differential expression across samples^10^.

Beside technical limitations, the development and complex structural organization of multicellular organisms itself complicates comparative gene expression analyses, as genes can have different expression patterns depending on tissue or life history stage^11^. The degree of specificity of a gene (i.e., the extent to which it is expressed broadly or restricted across conditions) has emerged as a biologically relevant parameter that is informative of function and robust for comparative analysis^10^. Gene expression specificity has proved useful when trying to understand the molecular basis of evolutionary constraints or tissue-selective pathogenesis^12,13^. The domain of expression of a gene is related to the pleiotropic effects that mutations in the gene might have. It can also support or contradict intuitions about core and specialized molecular functions and the definition of housekeeping genes^14–16^, as well as provide a frame of reference to analyze patterns of evolution of gene expression^10^.

Specificity is a relative attribute that depends on the context in which it is estimated. In the age of single-cell transcriptomics, the context in which expression specificity can be assessed is greatly expanded, opening new opportunities to study the molecular basis of organ and cell class variation, and how these patterns vary across species. Genes that might seem very specific because they are mainly expressed in cells of a given class can, at the same time, present varying degrees of specificity when considering cell subclasses or subtypes. This situation can be particularly relevant for highly specialized but abundant and heterogeneous cell classes. Neuronal cells, for example, are clearly distinct from other cell types but also very numerous and diverse within a given organism^17,18^.

Many methods have been proposed to estimate expression specificity^10^. The focus of common methods is to provide a score per gene that quantifies its level of specificity. This paper introduces an intuitive methodology to analyze gene specificity collectively by constructing global specificity landscapes. This global and comparative approach is achieved by first representing individual gene behavior graphically as a decreasing line relating expression levels with expression breadth -- i.e., an expression level-breadth (L-B) relationship. By compressing gene behavior in this way, we are able to easily quantify, globally visualize, and compare patterns of gene expression; which altogether provides a simple way to characterize and explore gene specificity patterns in different contexts. We use this methodology to characterize the specificity of human protein-coding genes individually and in functionally related groups across tissues, cell types, and species. We investigate the latent heterogeneity of neuronal-specific genes as an illustrative case of comparative analysis across contexts in the same organism. We study the evolutionary divergence of brain cell expression in 4 primate species to illustrate a comparative approach across organisms. All analytical tools are integrated in the software package *GeneSLand (*https://github.com/davilavelderrainlab/GeneSLand), together with precomputed specificity metrics and reference landscapes to facilitate future specificity analyses.

## Results

### Expression level and breadth are both relevant for specificity

A common approach to study expression specificity is by estimating the number of tissues in which a gene is expressed^12^. This parameter, often referred to as expression breath (B)^3,13,19^, provides an intuitive measure of expression specificity. A gene expressed by many tissues is not specific, whereas one expressed by a few is. But how do you decide if a gene is expressed or not? Expression breath is commonly estimated based on an arbitrary expression level over which a gene is considered to be expressed. The same cut-off value is used for all genes. Because the baseline expression level of a gene is very variable, conclusions such as how many tissues express certain kinds of genes could vary greatly depending on the chosen value. Consider as example the distribution of the number of tissues in which a gene is expressed. Estimating this distribution across 50 human tissues clearly shows a dependency on the chosen expression level threshold. At a value of expression greater than zero (TPM>0), the data is unimodal and left-long-tailed, suggesting that most genes are expressed in all tissues and very few, if any, are specific. At higher values (e.g., TPM>1, 2, or 3) the distribution starts displaying a U shape, suggesting a much larger fraction of specific genes (**Supplementary Fig. S1a**). General conclusions about the general specificity properties of genes will then depend on the threshold chosen. Expression level and expression breadth are both relevant parameters when describing gene expression specificity, and one hard threshold value to decide whether a gene is expressed or not might not always be a good approach.

### Analyzing gene specificity via L-B relationships

We propose an alternative, threshold-independent analysis that represents a gene’s expression pattern as a simple decreasing line describing the relationship between gene expression levels (x-axis) and gene expression breadth (y-axis) (**Fig. 1a-b**). Gene expression levels (L) are estimated by defining a certain number of bins (50 by default) covering the entire range of values in the dataset. The expression levels corresponding to the bins are used as increasing threshold values. Expression breadth (B) is estimated as the corresponding fraction of samples in which the gene is expressed at each threshold value. The shape of this L-B relationship line will then characterize the gene’s specificity behavior. If a gene is highly expressed in many samples, its L-B line will start at a high value in the y-axis and will show a long initial horizontal segment (**Fig. 1a-b**, gene A, red). If the expression value is very similar across the samples, the line will sharply drop at some L value (x-axis). It will decay gradually if it is variable (**Fig. 1a-b**, gene C, yellow). In the other extreme, if a gene is highly expressed only in one sample, its L-B line will first decay and then show a long horizontal segment until it drops to zero at a given L value (**Fig. 1a-b**, gene B, blue).

**Figure 1.**
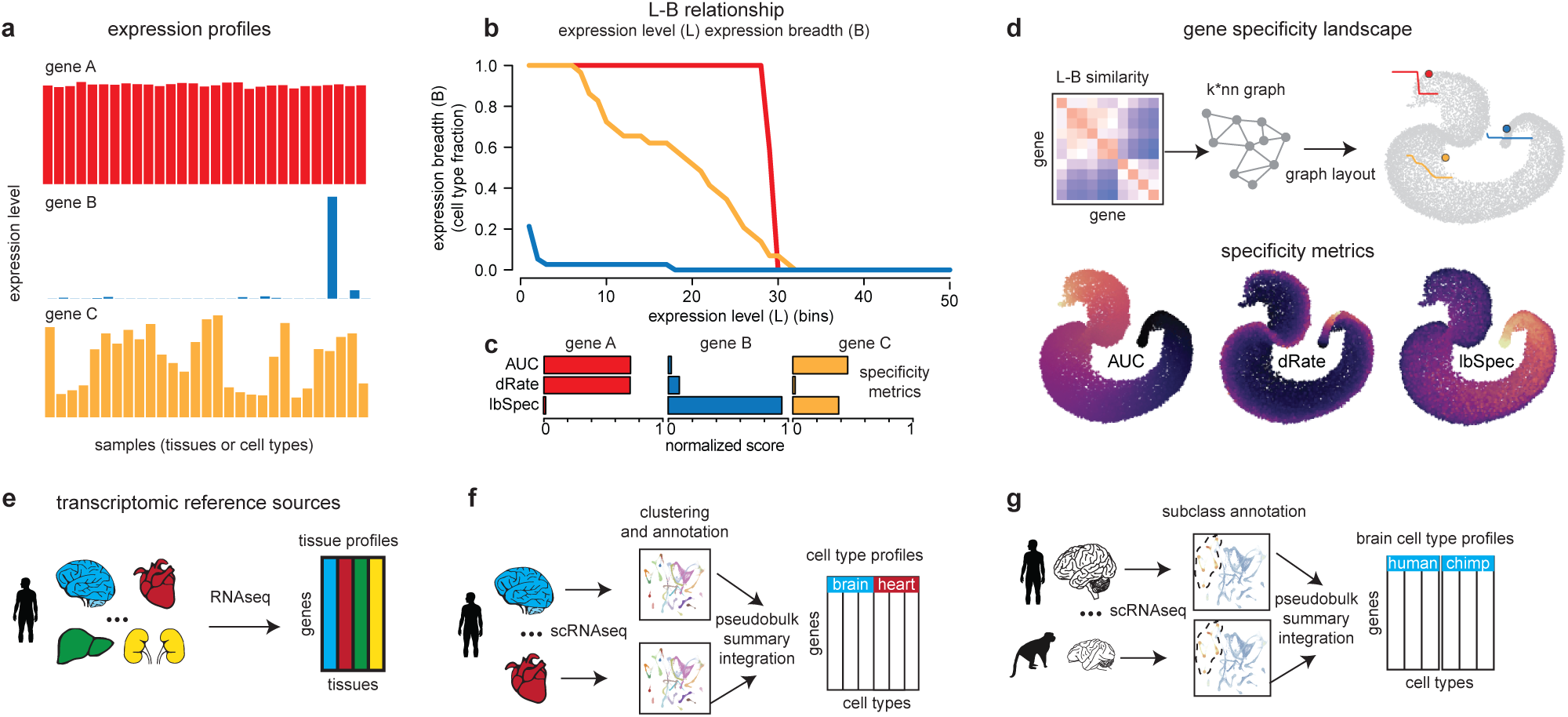
Gene specificity landscape framework. **a)** Expression levels of three representative genes across samples. **b)** Expression level (L; x-axis) vs expression breadth (B; y-axis) lines visualization of the three representative genes. Expression levels are divided into 50 expression bins. **c)** *AUC*, *dRate* and *lbSpec* scores of the three genes. Score max-normalization is computed over all genes. **d)** 2-dimensional landscape construction and score visualization. Scores are mapped to a color gradient from dark (low scores) to bright (high scores). **e-g)** Transcriptomic reference sources considered: human tissue profiles from bulk RNAseq data (e); human cell type pseudobulk profiles from single-cell RNAseq (scRNAseq) data (f); primate brain cell type profiles derived from scRNAseq data (g).

To quantitatively characterize the different behaviors, we introduce three metrics based on these lines. We use the area under the curve (*AUC*) as a measure of overall expression level and breadth. We use the rate of decay of the L-B relationship (*dRate*), estimated as the change in B over the change in L across variable B values, as a measure of expression homogeneity. Finally, we derived a new measure of specificity (*lbSpec*) based on the observation that highly specific genes will tend to have an initially low (B value), and broadly expressed genes will tend to display a sudden drop in the L-B line (**Supplementary Fig. S1b**). The *lbSpec* score quantifies the balance between these two behaviors across possible B values (Methods). A gene that is highly expressed at similar values across all or most samples will have high *AUC* and *dRate* values and a low *lbSpec* score (**Fig. 1c**, red). A gene that is highly expressed in only one or a few samples will have high *lbSpec* and low *AUC* and *dRate* values (**Fig. 1c**, blue). Genes with expression patterns between these extreme cases and varying expression levels across samples will tend to have half-maximum values of *AUC* and *lbSpec* scores and low *dRate* values (**Fig. 1c**, yellow). To simultaneously visualize, compare, and intuitively explore the specificity behavior of genes of interest in the context of all genes, we define a gene landscape based on L-B line similarity by building and projecting a gene-gene graph in 2D space. This landscape analysis gives us a globally consistent coordinate system in which we can project genes and their properties. Genes that localize close to each other in this landscape will have similar L-B behaviors and thus similar patterns of expression specificity (**Fig. 1d**).

Our analysis can be applied to any reference expression profiles, for example tissues, cell types, or subtypes (**Fig. 1e-g**). We illustrate the broad applicability of gene specificity landscapes by analyzing profiles at several levels of resolution: collections of bulk tissue samples (**Fig. 1e**), reference cell type profiles summarizing multi-tissue single-cell transcriptomic data (**Fig. 1f**), and summary profiles representing cell types and subtypes of the same organ (here brain) across several organisms (primates) (**Fig. 1g**). In what follows we apply our framework to these different contexts and demonstrate its robustness and consistency with biological expectations and alternative measures of specificity.

### Gene specificity landscapes capture constitutive and selective molecular processes

To develop intuition about the applicability of our approach, we start by analyzing reference transcriptomes of human tissues (**Fig. 2a**). The dataset includes 50 samples profiled with bulk RNA sequencing by the Human Protein Atlas (HPA) project. We estimated L-B lines for 18,848 genes, defined a gene tissue-specificity landscape, and calculated *AUC*, *lbSpec*, and *dRate* measures (**Fig. 2b**). Genes that are broadly and highly expressed across human tissues localized at one extreme of the landscape and tissue-specific genes at the opposite extreme. We observed a gradient from low to high *dRate* values across the whole landscape, indicating that genes can have very high or low variation in expression levels across tissues, irrespective of their overall expression level or specificity (**Fig. 2b**). Genes with extreme *AUC* or *lbSpec* values are consistent with biological expectations. The top 3 *AUC* values correspond to constitutively expressed mitochondrial genes, which display very high expression levels across all tissues (**Fig. 2c**) and corresponding high and long horizontal L-B lines (**Fig. 2d**). In contrast, extreme *lbSpec* genes include genes with very restricted expression (e.g., NOX3, MAGEC2), tissue-specific functions (e.g., SPO11 required for meiotic recombination in testis), and extremely low L-B lines (**Fig. 2c-d**).

**Figure 2.**
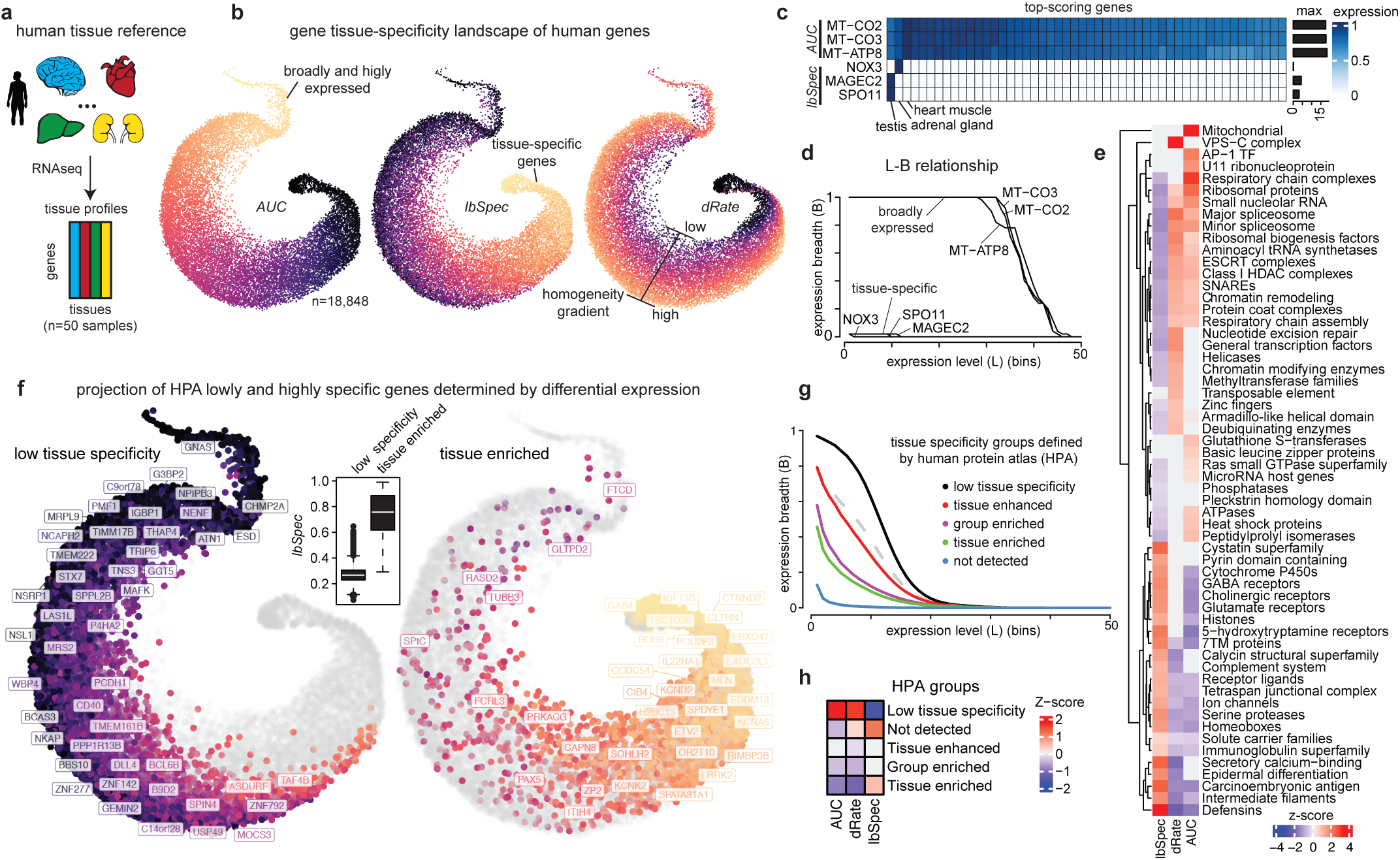
Gene tissue-specificity landscapes. **a)** Human tissue transcriptomic reference used as input. **b)** 2D gene specificity landscape of 18,848 genes. Data points represent individual genes. Scores are visualized using a color gradient from dark (low score) to bright (high score). **c)** Normalized expression of selected genes across tissues. The top three genes were chosen based on high *AUC*. The bottom three genes were chosen based on high *lbSpec*. Expression for each gene was divided by its maximum value. The right barplot shows the highest absolute expression per gene. **d)** Corresponding L-B lines of the selected genes. **e)** Enrichment analysis of specificity scores for HGCN gene groups with at least 10 genes. Top 10 groups per score are shown. **f)** 2D gene specificity landscapes highlighting low tissue specificity (left) or tissue enriched (right) genes as determined by the human protein atlas (HPA). Color gradient represents *lbSpec* scores. Boxplot insert shows *lbSpec* scores for the genes in the two groups. **g)** L-B lines of tissue specificity groups defined by the HPA. The dashed grey line represents random expectation. **h)** Enrichment analysis of L-B specificity metrics on HPA specificity groups.

To generalize these observations, we estimated *AUC*, *lbSpec*, and *dRate* values for reference gene groups from the human gene nomenclature committee (HGNC). We compared these estimates with expectations from randomly selected genes and identified gene groups with unexpectedly high values (enriched) (**Fig. 2e**). Consistent with gene-level observations, Mitochondrial and Respiratory chain genes showed the highest *AUC* enrichment, followed by ribosomal and RNA processing genes, as well as several other core constitutive processes (e.g., Chromatin remodeling, spliceosome, ATPases, heat shock proteins). Several of these constitutive processes also showed unexpectedly high *dRate* levels, indicating low variability across tissues in these highly expressed gene groups. In contrast, gene groups enriched with high *lbSpec* levels include highly specific receptors and sensory proteins, ion channels, and innate immune genes (e.g., defensins) which are not enriched (not different from random) or are depleted of *dRate* and *AUC* levels (**Fig. 2e**). Gene groups with a low, mid, or high-level tissue specificity are easily contrasted by computing average L-B lines and gene landscape projections, as illustrated by Ribosomal proteins, transcription factors (TF), and GABA receptors (**Fig. 2a-b**).

A common approach to define tissue-specific genes is by direct estimation of genewise expression level changes across tissues. Using this approach the HPA project has classified genes into different elevated expression categories. Tissue enriched genes are those having at least four-fold higher level in a tissue versus any other, while tissue enhanced are those with a similar increase but relative to tissue average (Methods). To develop further intuition, we contrasted our L-B based scores against these categories (**Fig. 2f-h**). As expected, genes with low tissue specificity localized at the constitutive extreme of the landscape and tissue enriched genes to the opposite extreme, with the latter having significantly higher *lbSpec* values (**Fig. 2f**). Average L-B lines for elevated expression categories clearly illustrate the different behaviors, with increasing specificity from low (black) to tissue enriched (green), with enhanced and group enriched (in related tissues) in the middle, and genes classified as not detected (very lowly expressed) in the opposite extreme (**Fig. 2g**). When compared with random expectation, “low specificity genes” showed enrichment of *AUC* values and tissue enriched enrichment of *lbSpec* (**Fig. 2g, Fig. S2c**). The latter was also enriched in “not detected genes”, indicating that this category includes those genes that are very specifically expressed in one or very few tissues at low levels. *dRate* values were enriched in both “low specificity” and “not detected” genes, indicating that both show a sharp drop in L-B lines, consistent with the broad gradient-like distribution of dRate values seen across the landscape.

Finally, we verified the robustness of the *AUC*, *lbSpec*, and *dRate* metrics to sample removal (Methods). In all cases, scores were highly robust to increasing removal of up to 50% samples (correlation >0.96) (**Fig. S2d**). *AUC* was the most robust measure, followed closely by *lbSpec* and *dRate*. *AUC* and *lbSpec* showed low variability across random replicates of a given removal percentage (correlation > 0.99 and 0.96), while *dRate* was slightly more variable (>=0.84 correlation). We also corroborated that our new specificity measure (*lbSpec*) closely matches the behavior of the widely used and robust metric *Tau*^10,20^, further supporting the applicability and robustness of L-B based specificity analyses (**Fig. S2e**). Taken together, these results demonstrate that gene specificity landscapes provide a robust and intuitive approach to explore and quantify patterns of gene expression specificity.

### The cell type specificity landscape of GO biological processes

In addition to contrasting genes, it is useful to directly analyse the behavior of groups of genes. Our results suggest that we can use aggregate LB-lines to contrast gene group specificity (**Fig. S2a-b**). We developed additional metrics for gene groups to generalize this group-based approach. We illustrate this by analysing expression specificity for 5,368 GO biological processes (gene groups) across a reference set of 81 human cell types (**Fig. 3a**). We study the specificity of a group of genes of interest by considering their average behavior. We contrast this behavior with randomly chosen genes and estimate statistical deviations from expectation and within group variability (**Fig. 3b**). All gene groups of interest are in this way individually mapped to L-B group relationships that can then be globally analyzed in a gene group specificity landscape (**Fig. 3b-c**). As an example, highly cell-type specific biological processes (e.g., cilium movement, chemical sensing) will localize at one extreme of this landscape that is characterized by high *lbSpec* scores and low *AUC* values. Constitutive processes (e.g., cytoplasmic translation, ribosomal assembly, mitochondrial electron transport) will localise at the opposite extreme with low *lbSpec* scores and high *AUC* values (**Fig. 3d**). Group level aggregate metrics clearly capture the average behavior of individual genes, as seen in the pattern of expression across cell types (**Fig. 3f, Fig. S3a-b**). Comparable random gene samples give us a reference to quantify how unexpected the specificity behavior of a given group of interest is (**Fig. S3c-d**). The expression breadth of highly (lowly) specific gene groups is significantly lower (higher) than randomly expected at most expression levels, a pattern easily seen when plotting deviation lines (**Fig. S3c**). Similarly, we can also quantify how unexpected the values of the *AUC*, *dRate*, and *lbSpec* L-B measures characterizing the L-B gene group behavior are (**Fig. S3d**). As expected, core constitutive biological processes (mitochondrial respiration, ribosomal assembly, translation) show significantly high *AUC* and low *lbSpec* values, while highly specific processes (cilia movement, chemical sensing) also show significant but opposite deviations.

**Figure 3.**
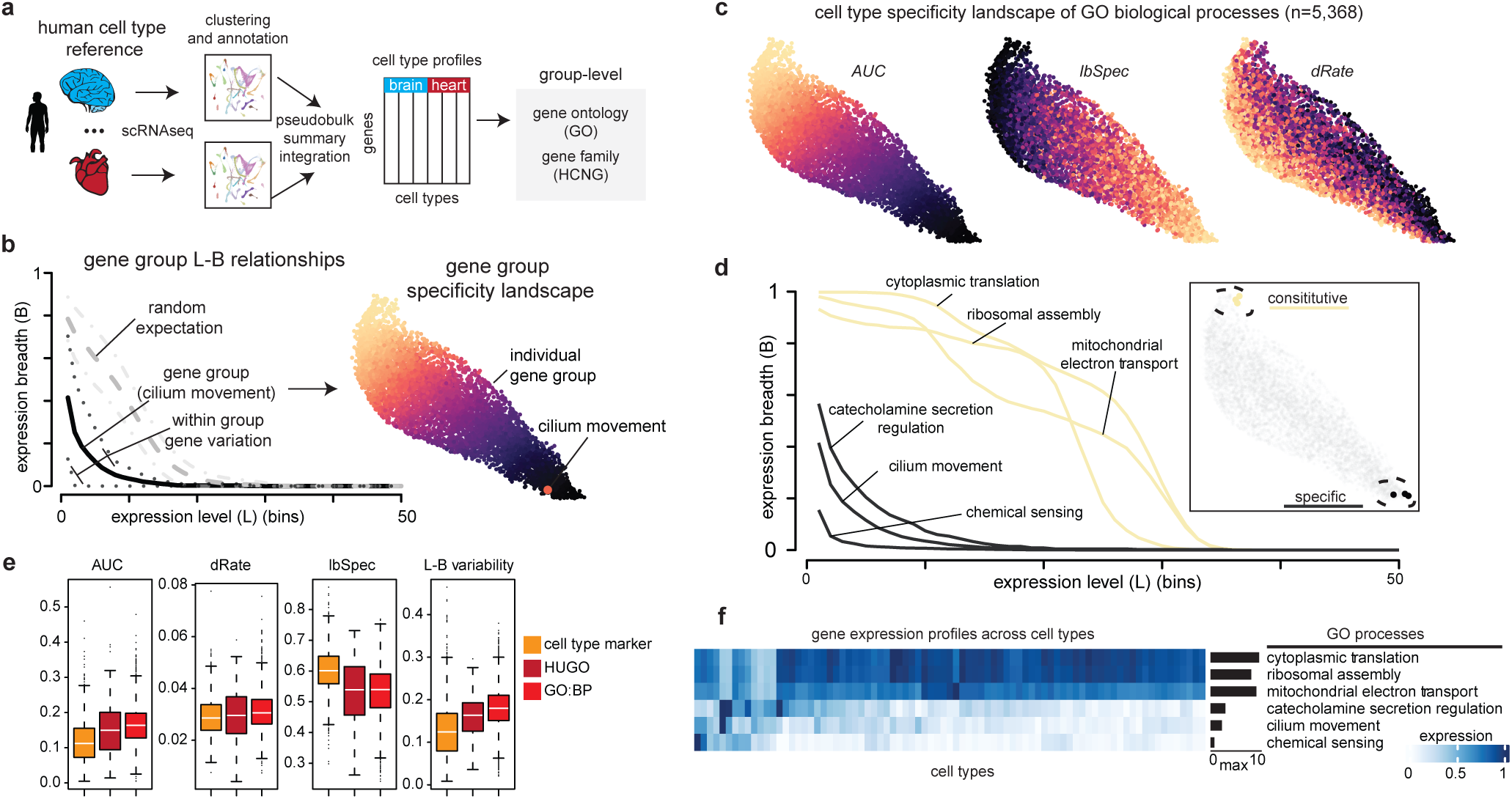
Cell type specificity landscape of GO biological processes. **a)** Human cell type transcriptomic reference used as input. **b)** Gene group L-B lines visualization. The black line represents the L-B scores computed on the mean expression of genes in a gene group. The highly specific “cilium movement” group is used as an example. The dashed black lines represent the mean plus or minus the standard deviation of the L-B scores computed over the individual genes in the group. The grey line represents the random expectation of gene groups with the same size as the analysed gene group. The dashed grey lines represent the mean plus and minus the standard deviation of genes in the random groups. The corresponding 2D gene group-cell type specificity landscape of 5,368 Gene Ontology biological processes is shown on the right. Each data point represents a gene group (GO:BP). Color gradient represents *AUC* scores. The “cilium movement” group is highlighted in the landscape. **c)** Cell-type specificity landscape of GO biological processes colored by L-B specificity scores -- color gradient from dark (low score) to bright (high score). **d)** L-B lines of three highly specific gene groups and three constitutive groups. Lines colored based on their *AUC* values (black: low *AUC*; yellow: high *AUC*). Insert on the right shows the corresponding location on the 2D landscape. **e)** L-B specificity scores (y-axis) distribution across gene categories. Each data point represents a gene group. L-B variability is defined across genes within a group. **f)** Normalized average expression of selected gene groups across cell types. Expression for each gene group was divided by its maximum value. The right barplot shows the highest absolute expression value per gene group.

Because the specificity of some gene groups is expected to be more variable than others, we tested whether our LB-based metrics capture such differences by contrasting groups of marker genes (cell type markers), gene families (HUGO groups), and biological processes (GO:BP). Marker genes are defined based on preferential expression in specific cell types and are thus expected to be specifically and similarly expressed. Gene families are defined based on homology relationships that often result in similar structure and partial molecular function conservation. Conserved function can be specific or constitutive, and thus families are expected to have low within group variability but not necessarily high specificity. Biological processes are defined based on common process involvement, regardless of molecular function or structure, and are thus expected to have high within group variability and mixed specificity. Our L-B group analysis of 1019 cell type marker sets, 144 gene families, and 5,368 GO biological processes confirmed these expectations, with higher specificity (*lbSpec*) and lower variability (L-B variability) in cell type markers than gene families and GO processes (**Fig. 3e**). Families showed intermediate variability and GO processes the highest. These results illustrate how L-B group level aggregate metrics and landscapes facilitate the intuitive comparative analysis of gene group specificity behavior.

### The specificity of neuronal genes

Genes that seem very specific because they are primarily expressed in cells of a given class can nonetheless present varying degrees of specificity when zooming in to consider only cells of that class. Gene specificity landscape analysis can help in contrasting these different contexts. We explored this situation of variable specificity in the context of neuronal cells as a proof of principle. Neurons form a well-defined and distinct class of cells that share morphological, functional, and molecular features^18^. At the same time, neurons are highly diverse, even within subclasses or types^21,22^. Molecular diversity within neurons has been shown to be primarily driven by genes of specific families^23^, many of these generally considered “neuronal”. We thus reasoned that genes identified as highly specific and neuronal relative to all cell types, would nonetheless display varying degrees of specificity when analyzing only neurons. We use this problem to demonstrate how to compare expression patterns across contexts via L-B analysis and specificity landscapes.

We first estimated gene-level L-B relationships across 81 human cell types to define a reference specificity landscape (**Fig. 4a-c**). To exemplify how highly specific genes distinctive of a class can seem broadly expressed when examining subclass specificity, we analyzed known pan neuronal and neuronal subclass genes. Neurexins, glutamatergic, and GABAergic genes are similarly expressed in neurons but not in other cell types and thus show highly specific L-B behavior across cell types (**Fig. 4a**). We include mitochondrial and GPCRs for reference. Mitochondrial genes are constitutively expressed and present broad L-B behavior. GPCRs are highly specific receptors and thus show L-B group behavior similar to that of neuronal genes (**Fig. 4a**). It would be easy to see these patterns by placing the genes in a reference landscape. To this end, it would be useful to fix a gene landscape and use it as reference, rather than recomputing one each time you have additional L-B lines. We thus developed a method to map L-B group behavior to the fixed 2D space of a reference landscape (Methods) (**Supplementary Fig. S4**). Mitochondrial genes and cell type specific neuronal and GPCR groups map at opposite extremes of the landscape, with the latter having high *lbSpec* values, as expected (**Fig. 4b**).

**Figure 4.**
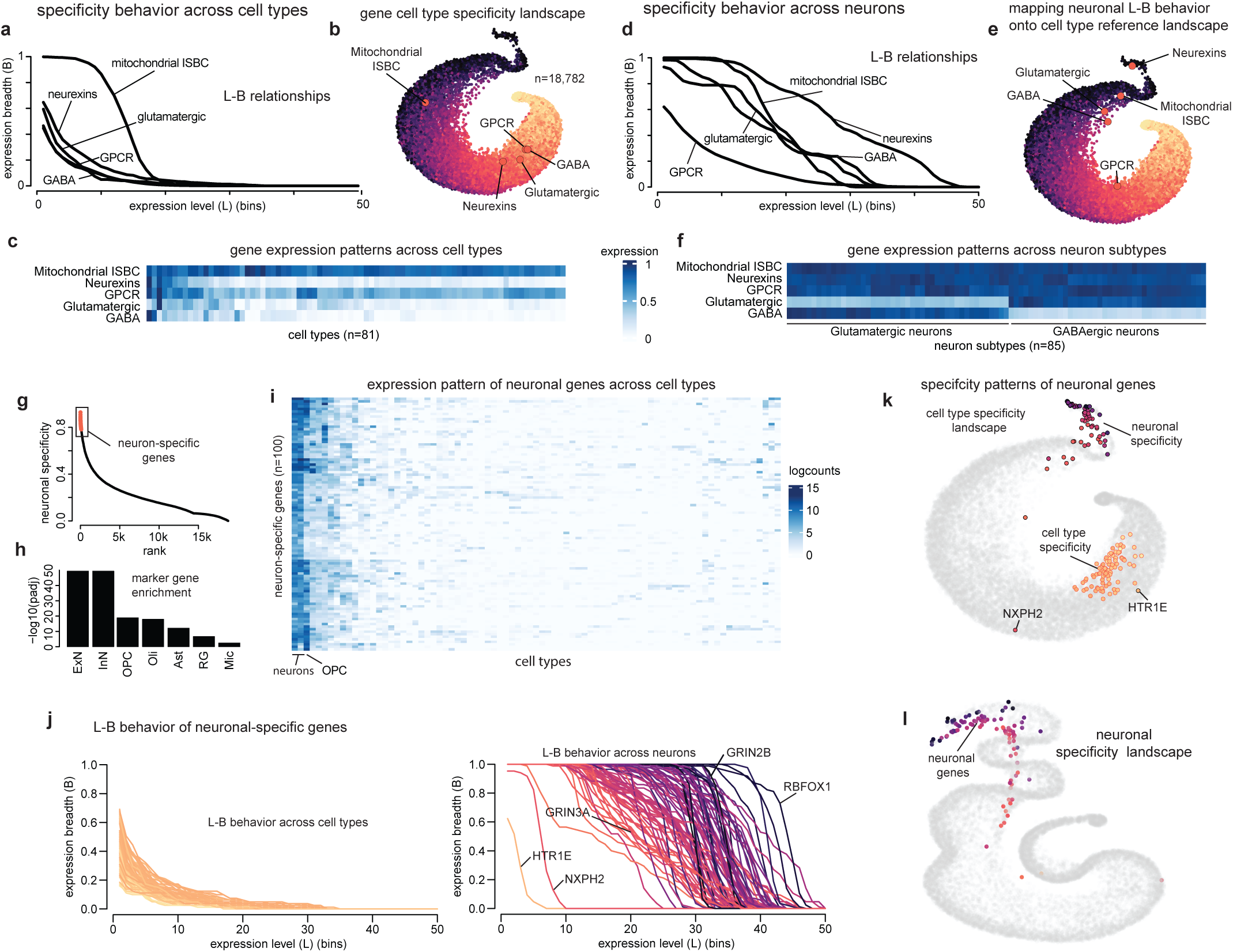
Expression specificity changes at increasing biological resolution. **a)** L-B lines of representative gene groups across human cell types. ISBC: Intermembrane Space Bridging Complex. GPCR: G-Protein Coupled Receptor. Groups were selected from HGCN groups. GABA and Glutamatergic groups were manually defined using marker genes. **b)** Cell type specificity landscape of 18,782 genes. The representative gene groups are mapped onto the 2D landscape based on their L-B behavior across human cell types. **c)** Normalized average expression of representative selected groups across human cell types. The expression for each gene group was divided by its maximum value. **d)** L-B lines of representative gene groups across human dorsolateral prefrontal cortex (dlPFC) neuron subtypes. **e)** Representative gene groups are mapped onto the 2D cell type specificity landscape based on their L-B behavior across dlPFC neuron subtypes. **f)** Normalized average expression of representative groups across dlPFC neuron subtypes. Expression for each gene group was divided by its maximum value -- Glutamatergic neurons subtypes on the left, GABAergic neuron subtypes on the right. **g)** Ranked neuronal specificity score. Red points highlight the top-100 genes**. h)** Enrichment of known marker genes in high-scoring neuronal specificity genes (p-value, GSEA rank-based analysis). **i)** Expression pattern of top-100 neuronal specific genes across reference cell type profiles (HPA). **j)** L-B behavior of the top-100 neuronal specific genes. On the left, L-B behavior in reference cell types. On the right, L-B behavior across neuron subtypes. **k)** Neuronal specific genes mapped onto reference cell type reference landscape. Black circled dots represent neuronal specific gene behavior across neuron subtypes. Red circled dots represent neuronal specific gene behavior across reference cell types. The color of the dot represents specificity values, with lighter color indicating higher specificity. **l)** Neuronal specific genes in the neuron subtype landscape. The color of the dot represents specificity values, with lighter color indicating higher specificity.

We next zoom in to explore neuronal specificity. We estimated gene-level L-B relationships across 85 neuronal subtypes from the human prefrontal cortex (Methods) (**Fig. 4d-f**). Neurexins are cell surface molecules expected to be expressed in most neurons, while glutamatergic and GABAergic genes are discriminatory of neuronal subclasses and thus expected to be expressed in only a subset of neurons. Expression profiles are consistent with these expectations (**Fig. 4f**). Neurexins show a broad L-B behavior, and glutamatergic and GABAergic L-B lines show mid specificity (in between Neurexins/Mitochondrial and GPCRs) when analyzing neuronal specificity (**Fig. 4d**). This change in specificity is reflected in a change in position when mapping the gene sets onto the fixed cell type specificity landscape (**Fig. 4e**). Thus, changes in L-B behavior and landscape position can help contextualize gene specificity.

We generalized this analysis to highly specific neuronal genes. We ranked genes based on neuronal specificity by considering their general cell type specificity (*lbSpec*) and their neuronal expression, both estimated from cell type reference data (Methods) (**Fig. 4g**). We next focused on the top-100 most neuronal specific genes. These genes have high *lbSpec* values and high expression levels in neurons. Overrepresentation of known neuronal marker genes (ExN, InN) (**Fig. 4h**) and expression profiles confirmed neuronal specificity (**Fig. 4i**). We then investigated their specificity within the context of only neuronal cells. As expected, their L-B behavior is highly specific across cell types (**Fig. 4i, left**). In sharp contrast, the L-B behavior of these neuronal-specific genes is heterogeneous across neuronal subtypes (**Fig. 4i, right**). Although seemingly equally specific at the level of cell types, our neuronal analysis uncovered varying levels of specificity. In one extreme, the neuronal GPCR serotonin receptor HTR1E and the neurexophilin NXPH2 (a putative signaling neuropeptide encoding gene) are highly specific, suggesting preferential activity in very specific neuronal subpopulations. Remarkably, even stereotypical neuronal genes such as NMDA glutamate receptors showed heterogeneity (i.e., a more specific subunit GRIN3A and a more broadly expressed GRIN2B), suggesting preferential subunit expression across neuronal subtypes. In the opposite extreme, we identified the neuron-specific splicing factor RBFOX1 as broadly and highly expressed across neuron subtypes (**Fig. 4j)**. These contrasting patterns of specificity relative to all cell types (bottom) or only to neuron subtypes (top) can be clearly appreciated when projecting the 2 contrasting L-B behaviors to the same reference landscape (**Fig. 4k**). The varying degrees of specificity of these highly specific neuronal genes are also apparent when exploring a gene specificity landscape constructed only for neuronal cells (**Fig. 4l**). Taken together, these results demonstrate how L-B analysis and specificity landscapes offer a simple way to explore and compare expression patterns across cellular contexts.

### Cross-species analysis of brain cell specificity

Genes are expressed at different levels at different times and in different tissues in multicellular organisms. It is thus more difficult to understand how gene expression parameters relate to molecular evolution in these organisms than in bacteria or yeast^11^. It has been proposed that robust measurements of expression specificity could be useful in studying the evolution of gene expression^10^. Comparisons between human and mouse provided a first support for tissue specificity as a strongly conserved and biologically relevant parameter^10^. We reasoned that we could use our L-B framework to further explore such conservation in the context of primates and brain cells. We obtained single-nucleus transcriptomic data profiling the prefrontal cortex of adult humans, chimpanzees, rhesus macaques, and common marmosets^24^. We estimated summary expression profiles for 92 subtypes of excitatory neurons (ExN), inhibitory neurons (InN), or glial cells (Glia) in each species; and sought out to investigate the conservation of gene specificity across species and cell classes (**Fig. 5a**). L-B lines provide a simple qualitative approach to explore this. We described the pattern of specificity of each species globally by the average L-B behavior of all genes and estimated how this behavior in other primate species deviates from that of humans. Remarkably, we found that these deviations mirror evolutionary divergence times, with chimpanzees (6.40MYA divergence from human) having the smallest deviation, followed by rhesus macaques (28.82MYA) and marmosets (42.9MYA) (**Fig. 5b**); suggesting that global changes in specificity capture evolutionary trends or transcriptome variation. The same qualitative pattern was observed irrespective of cell class, but the magnitude of deviation varied. The specificity of glia cells seems to have diverged the most, and inhibitory neurons the least. This pattern is primarily observed when comparing humans to chimpanzees, suggesting that gene expression levels and specificity among inhibitory neuron subtypes have remained relatively more constrained than those in excitatory neurons and glial cells since the human-chimp divergence. These global qualitative observations are consistent with direct measures of similarity of specificity profiles (*lbSpec*). In all cases genome-wide cell subtype specificity was highly correlated (Pearson’s correlation > 0.8), supporting the strong conservation of this expression parameter within brain cells (**Fig. 5c**). The strength of correlation again mirrored evolutionary divergence in all cell classes, further supporting the biological relevance of expression specificity. Finally, we show that our L-B calculations can be used for more in-depth gene-level analyses of expression evolution. By comparing specificity gene metrics across species, it is possible to identify individual genes with an apparent increase in expression levels and breadth across neurons during primate evolution (e.g., the Apolipoprotein A1 gene), as well as those whose expression has become more specific (e.g., the Carbonic Anhydrase 9 gene). These analytical tools and precomputed parameters thus open opportunities for systematic studies of gene expression changes in brain cells during primate evolution. All precomputed profiles, L-B landscapes, and metrics are included in Supplementary data and/or accompanying R package.

**Figure 5.**
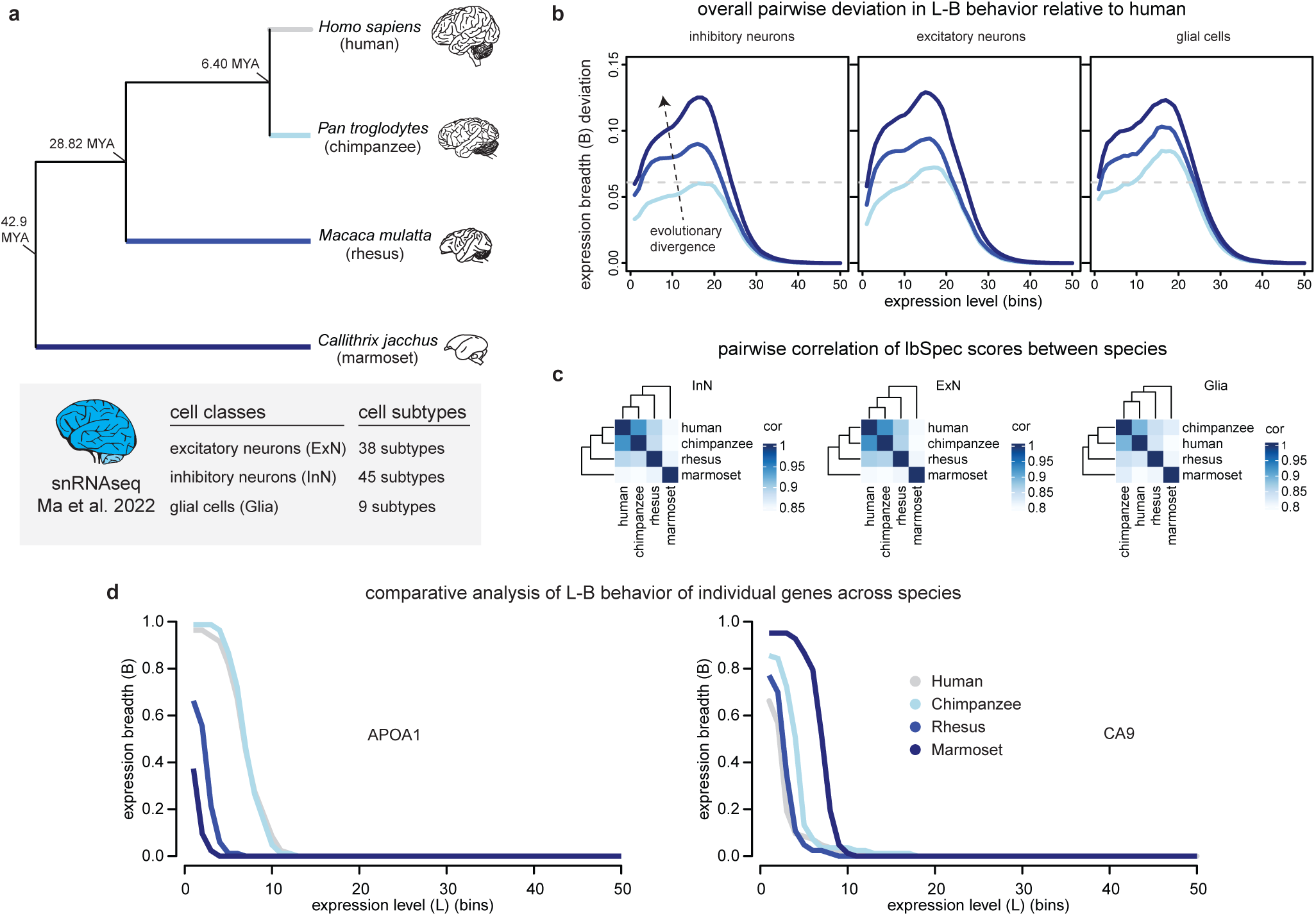
Deviations in expression specificity among primates. **a)** Phylogenetic tree of the four primate species considered (top). MYA: million years ago. Tree and divergence times were obtained from TimeTree (https://timetree.org/)^43^. Darker colors indicate higher evolutionary distance to humans (shown in grey). Description of snRNAseq data source (bottom). **b)** Mean absolute expression breadth (B) deviation between humans and each other primate species. The horizontal dotted line indicates the highest B deviation of chimpanzees in inhibitory neurons and provides a reference point. **c)** Pairwise correlation of *lbSpec* scores between primates. A darker color indicates a higher correlation. The correlations were computed separately for inhibitory neurons (left), excitatory neurons (middle) and glial cells (right). **d)** L-B lines of representative genes with changing specificity behavior across species. L-B analysis was performed across neuronal subtypes.

## Discussion

Gene expression specificity is a biologically meaningful parameter that is useful in understanding molecular evolution and disease etiology in multicellular organisms^3,12^. We present a novel methodology to analyze gene specificity by defining simple L-B relationships and specificity landscapes. We show that this approach allows qualitative analysis, quantification, visualization, and comparison of patterns of gene expression specificity in different contexts. We explore gene and gene group specificity within cell classes and across tissues, cell types, and species as different biological contexts.

Specificity is a very intuitive concept. However, its operationalization with quantitative data is not so simple. The simple approach of counting the number of tissues in which a gene is expressed is problematic. It depends on choosing an expression threshold that might not generalize and influence conclusions^10^. This motivated us to devise an approach that bypasses the need of selecting a cut-off value. We came up with the idea of representing expression patterns as L-B lines and realized that this approach also facilitates global comparative analyses and visualization. Thus, although many methods already exist to calculate a specificity score per gene^10,20,25–29^, our approach focused on analyzing gene specificity collectively by constructing global landscapes that facilitate qualitative, quantitative, and comparative analyses at the gene and gene group levels. L-B lines and reference landscapes provide an easy way to qualitatively explore how genes behave relative to other genes in the same context, as well as how their behavior changes in different contexts. L-B relationships map to quantitative parameters that numerically describe and compare the qualitative patterns. Landscapes provide a globally consistent coordinate system in which genes of interest and their properties can be projected. Like L-B derived metrics, the widely used method Tau^20^ does not depend on a cut-off value and has been shown to be the most robust among commonly used specificity methods^10^. The high consistency of this method with our *lbSpec* metric is reassuring and further supports its robustness and generalization. Our own resampling tests confirmed such robustness (**Fig. S2)**.

Although our L-B approach does not rely on an individual cut-off value, it does rely on a finite set of threshold values and a finite set of B values -- the latter determined by the number of samples. Therefore, a potential limitation is its applicability on datasets with a few samples. In such cases, many genes may have the same score in some of the metrics, limiting comparative analyses. A similar limiting situation would be one in which data quality leads to noisy profiles with many genes captured only in few samples and with low values. These genes will be scored with high specificity and thus the analysis would provide biased conclusions. The framework is designed for the analysis of high-quality and extensive reference datasets like those analysed here and included in our package.

It has been pointed out that the proportion of closely related samples (e.g., different brain regions) in the dataset under analysis might bias specificity measures^10^. Related to this aspect, in the current study we consider the addition of expression profiles derived from single-cell data to investigate the effects on specificity of closely related samples. We interpret this as an opportunity to dissect further the function of genes, and not exclusively a potential technical artifact to be aware of. As an example, the specificity of certain kinds of genes across neurons of a given type can be informative of the molecular basis of their function^23^. We showed how this kind of problem can be explored by changing the context in which specificity is estimated. L-B lines and landscapes can be useful in “zooming in” analyses of this kind. This can be useful in additional contexts where cells of a class share core properties but also have specific adaptations, for example tissue-resident immune cells^30^. Single-cell biology opens many opportunities for the in-depth study of cellular specificity. We consider our landscape approach can provide an analytical platform to facilitate those analyses.

Gene specificity is also a biologically relevant parameter for studying the evolution of gene expression. Similar to the overall conservation of tissue specificity between orthologous genes in mouse and human^10^, using our analysis framework we found that brain cell specificity is highly conserved across primates. By including 4 primate species, we found that the level of conservation of specificity patterns within brain cell classes mirrors evolutionary divergence times. This result is consistent with previous gene expression divergence estimates based on correlation and differential expression^24^. The larger divergence of glial cells in primates is also supported by such estimates as well as comparative transcriptomic studies at the level of bulk tissue and cell-sorted populations^31,32^. Comparing expression levels between species is not trivial^33,34^. Our results suggest that L-B lines and their quantitative measures can be one way to operationalize evolutionary comparisons within cell and evolutionary lineages. More in-depth analyses are required to systematically identify and validate specific expression changes in similar neuronal subtypes across species. We hope the reference landscapes and measures reported here could provide raw material to identify promising candidates for follow-up investigation.

In conclusion, gene specificity landscapes provide a robust and intuitive approach to explore and quantify patterns of gene and gene group expression specificity. Beyond introducing additional specificity metrics, our framework provides a means to generate reference platforms to explore biological questions of interest. We generated L-B relationships, landscapes, and measurements for human genes and gene sets across tissues, cell types, and brain cells. All analytical tools and resources in the R package *GenesLand* are provided to promote such explorations.

## Supplementary Figures

**Supplementary Figure 1.**
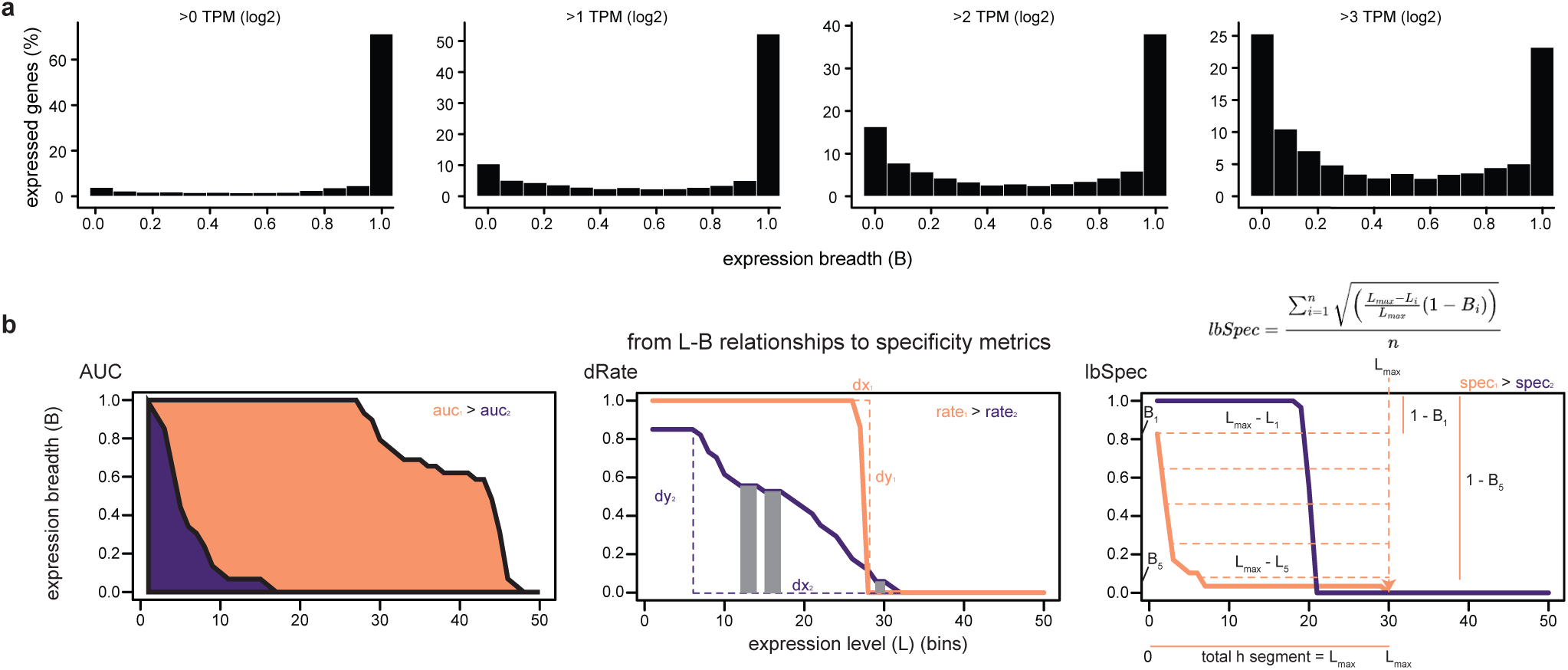
Expression breadth distribution and L-B specificity metrics. **a)** Percentage of expressed genes (y-axis) having a given expression breadth (x-axis). From left to right, an increasing TPM threshold is considered to define a gene as expressed or not. **b)** Schematic representation of *AUC*, *dRate*, and *lbSpec* metric calculation. Purple lines represent a low scoring gene and orange lines a high scoring gene. *AUC* (left): colored areas represent the area under a curve (*AUC*). *dRate* (middle): grey areas represent portions of the line with a constant expression level. These portions are not considered in the computation of the *dRate* score. Dashed lines represent the variation in expression level or expression breadth. *lbSpec* (right): horizontal dashed lines represent the expression level segment considered for each expression breadth value. Vertical dashed lines represent the expression level at which the expression breadth reaches zero.

**Supplementary Figure 2.**
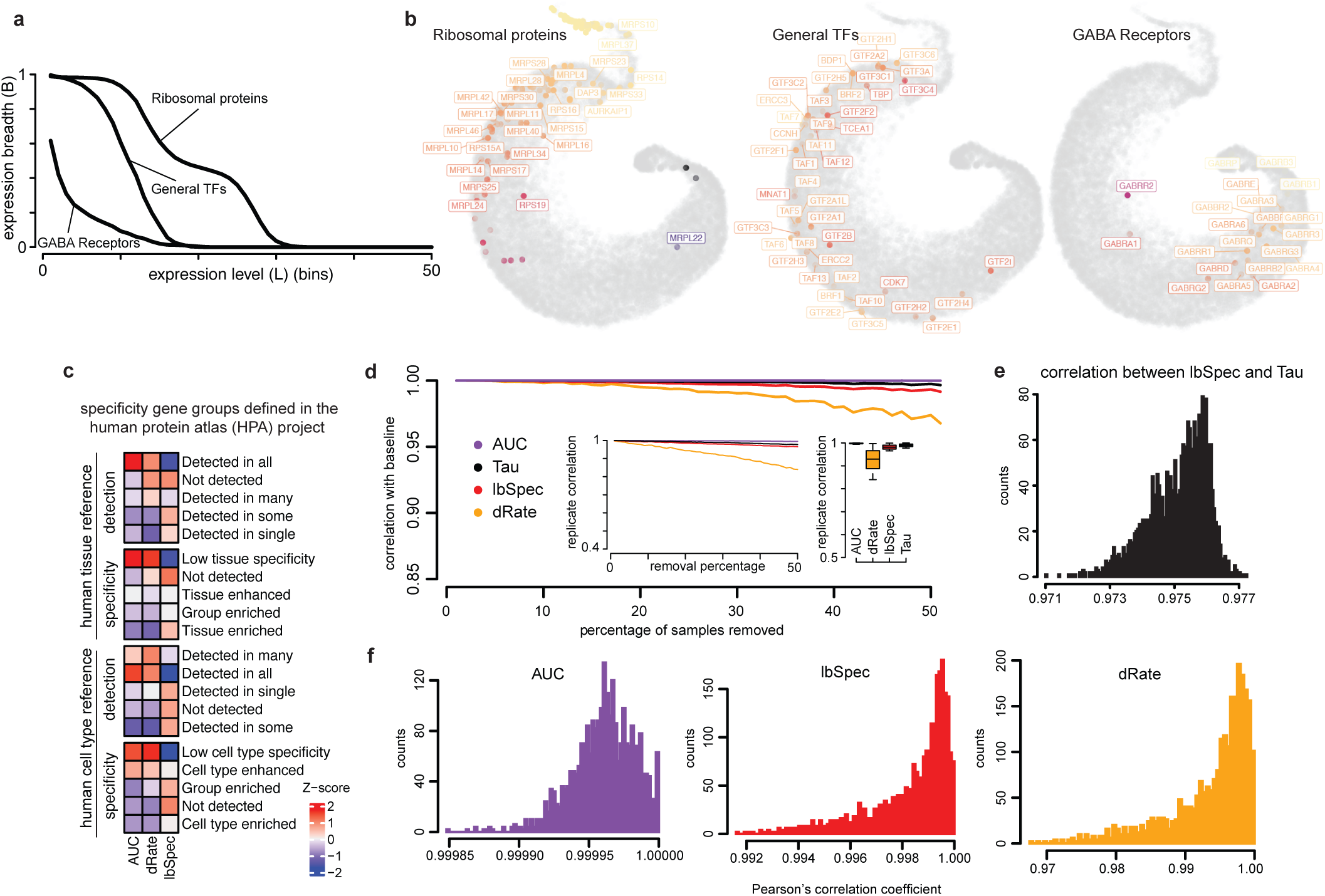
L-B behavior of gene groups and robustness analyses. **a)** L-B lines for ribosomal proteins (high), transcription factors (general TFs) (middle) and gamma-aminobutyric acid (GABA) receptors (low). **b)** 2D gene tissue specificity landscapes highlighting genes belonging to ribosomal protein (left), general TF (middle), or GABA receptor (right) groups. Genes are colored based on the *AUC* score. **c)** Enrichment analysis of L-B specificity metrics for HPA specificity and detection gene categories defined across tissues (above) or cell types (below). **d)** Correlation between baseline score values and values after sample removal (x-axis). *AUC* is shown in purple, *Tau* specificity in black, *lbSpec* in red, and *dRate* in yellow. The line plot insert shows the average correlation between random removal replicates per removal fraction (%). The boxplot insert shows random removal replicate correlation across all sample removal percentages. **e)** Distribution of correlation values between *lbSpec* and *Tau* metric across all replicates and sample removal experiments. **f)** Distribution of correlation values across sample removal percentages.

**Supplementary Figure 3.**
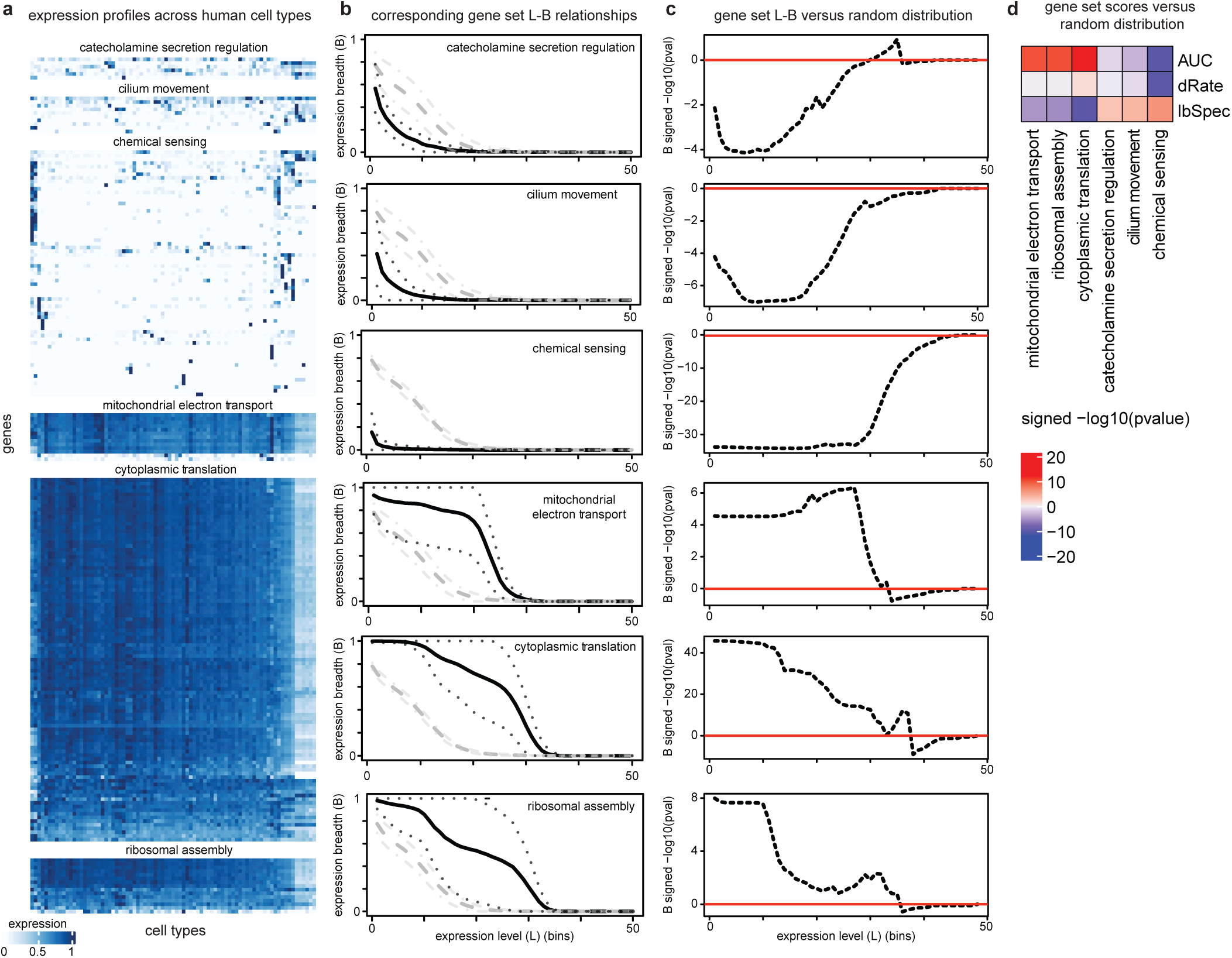
L-B behavior of specific and constitutive processes. **a)** Normalized expression profiles across human cell types for each gene of representative specific (top) and constitutive (bottom) gene groups. GO biological processes were selected based on extreme *lbSpec* (top three) or *AUC* (bottom three) values. Gene expression was divided by its maximum value. **b)** L-B lines of representative gene groups (black), along with the intergenic L-B variability (dashed black), the average random expectation (grey), and the variability of the random expectation (dashed grey). **c)** Deviation from random expectation of L-B behavior. Z-scores were computed individually by expression level bin. Data points represent −log(pvals) multiplied by 1 in case of positive deviation (enrichment, z-score>0) or −1 for negative deviation (depletion, z-score<1). The red line indicates a value of zero. Lines that for the most part show negative (positive) values indicate unexpectedly low (high) expression breath values and thus high (low) specificity. **d)** Enrichment analysis of L-B specificity metrics for representative gene sets.

**Supplementary Figure 4.**
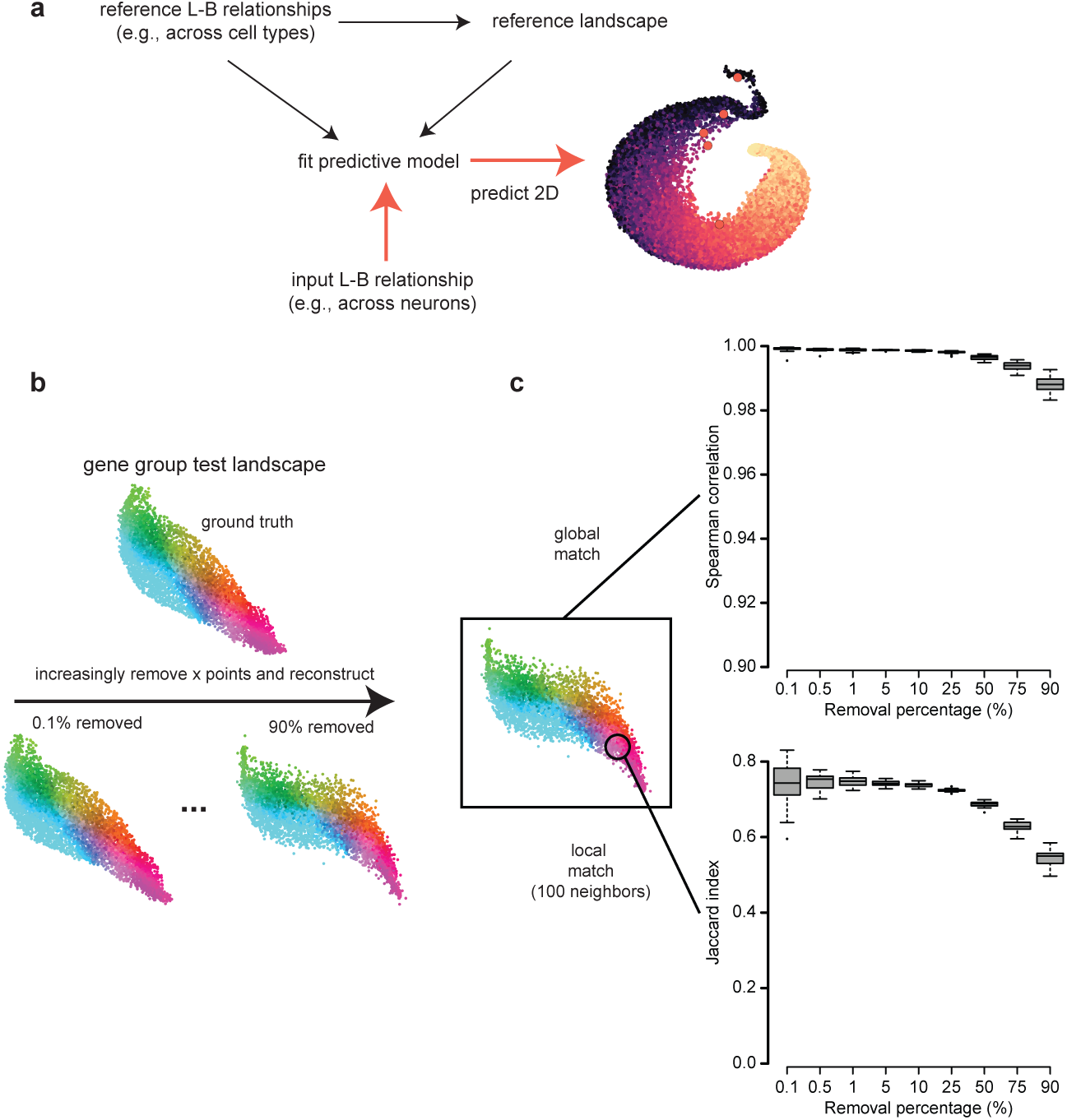
Reference landscape mapping procedure. **a)** Procedure to map input L-B relationships to a precomputed reference landscape. **b)** Evaluation of the mapping procedure. **c)** Global match scores between predicted and reference landscape positions (top). Global match was measured by the consistency of the relative location between predicted and reference positions as measured by the rank correlation of distance vectors (Methods). Match scores between predicted and reference local neighborhoods (bottom). The match is measured by the overlap or top-100 neighbors and quantified by the Jaccard index (Methods).

## Methods

### Transcriptomic data

#### Tissue and cell type references

Transcriptomic profiles for human reference cell types (n=81) were downloaded from the Human Protein Atlas (HPA) project (https://www.proteinatlas.org/about/download). Reference transcriptomes are based on the meta-analysis of single-cell RNA sequencing (scRNAseq) data from 31 human tissues as described in (https://www.proteinatlas.org/about/assays+annotation#singlecell_rna) and reported in^35^. Pseudobulk profiles summarizing cell type clusters and normalized in transcript per million (TPM) values per protein-coding gene were used as reference cell type profiles (18,782 genes, *rna_single_cell_type.tsv*). Consensus transcriptomic profiles for human reference tissues (n=50) were also downloaded from the HPA project. Normalized expression (nTPM) values calculated as the maximum TPM value for each gene from HPA and GTEx data were used as tissue reference profiles (18,848 genes, *rna_tissue_consensus.tsv*). The data is based on The Human Protein Atlas version 23.0 and Ensembl version 109. <u>Brain cell type references</u> scRNAseq data of the dorsolateral prefrontal cortex (dlPFC) from 4 primate species (human, chimpanzee, macaque, marmoset) reported in^24^ were downloaded from http://resources.sestanlab.org/PFC/. The dataset includes expression data for 172,120 single cells and 16,003 protein coding genes. Pseudobulk summary profiles were computed for 113 cell subtypes by summing gene counts and then normalizing to obtain counts per million (CPM). All pseudobulk profiles used in our analyses are reported in **Supplementary data**.

### Gene groups

#### HGCN

Gene groups defined by the Human Gene Nomenclature Committee (HGCN) were downloaded from https://www.genenames.org/download/gene-groups/%23!/%23tocAnchor-1-2. Only parent terms (terms that do not have any parent node) were considered for analyses. <u>CellMarkers</u> Human cell markers were obtained from CellMarker 2.0^36^. Only non-cancer cells were considered. <u>Gene Ontology Biological Processes</u> Gene Ontology Biological Processes (GO:BP)^37,38^ were downloaded from the Enrichr website (https://maayanlab.cloud/Enrichr/#libraries) (GO_Biological_Process_2025). <u>GABA and Glutamatergic</u> The GABA set was defined by the following three genes: GAD1, GAD2, SLC32A1; the Glutamatergic set by the following three: SLC17A7, SATB2, TBR1. When calculating gene group behavior, only gene sets with transcriptomic data for at least 2 genes were considered for analysis.

### HPA gene expression classification

Expression specificity groups defined by the HPA were downloaded from the Human Protein Atlas version 23.0 (https://www.proteinatlas.org/about/download, *proteinatlas.tsv*). Specificity categories were defined as follows: “tissue enriched genes” have at least fourfold higher mRNA levels in one tissue compared to all other tissues; “group enriched genes” have at least fourfold higher mRNA levels in a group of 2–5 tissues compared to all other tissues; “tissue enhanced genes” have at least fourfold higher mRNA levels in one tissue compared to the average level in all other tissues; “low tissue specificity genes” have mRNA levels above cutoff in at least one tissue but do not belong to any of the above categories; “not detected genes” have mRNA levels below cutoff in all tissues. Distribution categories are defined as follows: “genes detected in single” are genes for which a single tissue has detectable levels; “detected in some” are genes where more than one but less than one-third of the tissues have detectable levels; “genes detected in many” have detectable levels in at least a third but not all tissues; “detected in all” are genes where all tissues have detectable levels; “not detected” are genes where none of the tissues have detectable levels^39^.

### Expression level-breadth (L-B) analysis

A gene’s expression pattern is represented as a decreasing line describing the relationship between gene expression levels (x-axis) and gene expression breadth (y-axis). Gene expression levels (L) are estimated by defining a certain number of bins (50 by default) covering the entire range of expression values in the dataset. The expression levels corresponding to the bins are used as increasing threshold values. Expression breadth (B) is estimated as the corresponding fraction of samples in which the gene is expressed at each threshold value.

### Expression specificity metrics

#### *AUC* score

The Area Under the Curve (*AUC*) of an L-B line is computed following the trapezoid rule.

#### dRate score

The decreasing Rate (*dRate*) is computed numerically as the average change in B with respect to L across the L-B line considering only nonzero values.

#### lbSpec score

L-B lines of broadly expressed and highly specific genes are qualitatively distinct. Genes that are highly expressed in many samples will have a long top-left horizontal segment close to a value of 1 in the y-axis. Those expressed highly in few samples will have a horizontal segment that is long at the bottom-right close to a value of 0 (**Fig. S1b**). A specificity index *lbSpec* was formulated to capture this intuition and quantify the degree to which any L-B line approximates those extreme behaviors. The index *lbSpec* takes values between 0 and 1, with 0 corresponding to highly and broadly expressed and 1 to highly specific genes. It is computed as follows:

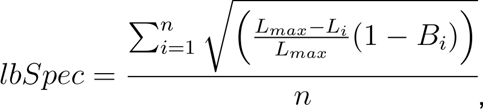

where *L_max_* is the value of L where the L-B line reaches a value of B=0. The *L_max_* thus corresponds to the maximum expression level at which the gene is expressed in at least one sample and defines the size of a reference total horizontal segment for the gene (**Fig. S1b**). *L_i_* and *B_i_* are the set of L and B values spanning the L-B line. The *lbSpec* index is thus a weighted average of the horizontal segments covered by the line at the different expression breadth values (B). The weight 1 - *B_i_* represents the proportion of samples in which the gene is expressed at lower values than the corresponding *L_i_* value and ensures that lines with long top-left horizontal segments (broad expression) will have low *lbSpec* values and lines with long bottom-right horizontal segments (highly specific) will have high *lbSpec* values. A graphical illustration of this calculation is presented in **Fig. S1b.**

### Gene specificity landscape construction

A global gene specificity landscape is built by connecting genes based on the similarity of their L-B behavior and projecting the resulting graph into 2D space using a graph-layout algorithm. The gene-gene graph is built using a density-dependent nearest-neighbor graph (k*NN) algorithm^40^. Similarity is measured using Euclidean distance as a metric. A modified version of the stochastic gradient descent (SGD)-based layout algorithm UMAP is used as graph-layout algorithm (the one used by UMAP) taking as input the k*NN network and per-gene initial positions. Initial positions are defined by approximate right singular vectors computed by randomized singular value decomposition (SVD) as implemented in R package *irlba*. Landscapes for gene groups are built by first computing L-B lines based on the mean expression of genes in each gene set. Graph construction and layout are performed as implemented in the *buildNetwork* and *layoutNetwork* functions of the R package *ACTIONet*^41^.

### Specificity Single Cell Experiment object

A Specificity Single Cell Experiment (SSCE) object is used as data structure to organize the L-B and landscape analyses. The object is a simple adaptation of the standard Single Cell Experiment (sce) object^42^ to encode the L-B lines of a gene or gene set as a 50-dimensional column vector within an assay matrix of the sce object (dimension 50 x number of genes/gene sets). *AUC*, *dRate*, and *lbSpec* scores are stored as column metadata in the same object (*colData*), along with initial B values (*initB*), x and y landscape coordinates, and a color palette.

### Statistical analysis of group behavior

Deviation of the L-B behavior of a gene set from random expectation is estimated by comparison with 200 random sets with the same number of genes and quantified using a z-score and corresponding p-value. Estimates are computed for each metric (*AUC*, *dRate*, and *lbSpec*). The variability of a gene set is measured by the standard deviation across the genes within the set. Variability is estimated for L-B lines (each individual B value and their average) and for specificity measurements (*AUC*, *dRate*, and *lbSpec*). Variability measurements are annotated in SSCE metadata.

### Robustness analysis

The robustness of the *AUC*, *lbSpec*, and *dRate* measures was assessed by estimating the effect of random sample removal. Samples were randomly removed at percentual increments from 1 to 50%. Random removal and estimation were performed 20 times per removal step. The effect of removal was assessed for both equal-size removal replicates and across replicates during progressive removal using the Pearson’s correlation coefficient.

### Landscape mapping

Mapping of new genes or gene sets to a pre-existing reference 2D landscape is performed using a predictive model that takes as input L-B lines (ordered B values) and predicts corresponding x, y coordinates (**Fig. S4a**). The model is first trained to learn the mapping from L-B lines to 2D coordinates of the reference landscape. The random forest algorithm implemented in the R package *ranger* was used for all predictions. Predictions for input L-B lines are performed using the trained model and the R package *stats*. The predicted coordinates are then used to map the new genes or gene sets over the landscape for visualization. <u>Evaluation</u> This mapping procedure was evaluated by measuring the degree to which the predicted local and global positions of test gene sets in a reconstructed landscape recover their known position in a reference landscape (**Fig. S4b**). For this evaluation, the cell type specificity landscape of GO biological processes was used as reference. The procedure was conducted as follows. Consider a fraction *x* of GO biological processes. These processes are removed from the data and a new reconstructed landscape that does not contain the x processes is built. The corresponding position in the reconstructed landscape (2D coordinates) is then predicted for the missing processes using the remaining data for training. The goal of the evaluation is to assess how well the predicted position matches the position of the same processes in the complete reference landscape. This was assessed using two complementary approaches. A global approach assesses the match of the location of the *x* processes relative to the position of all other processes. To quantify this match, first the Euclidean distance of a predicted point to all other points in the landscape is used as an estimate of relative position. This estimate is computed for all *x* processes in both the reconstructed and the reference landscapes. Then the match between the relative positions in both landscapes is quantified by rank correlation. If a point *x* is close and distant to similar processes in both landscapes the match is high and the correlation value close to 1. A second, local approach assesses the match between the local neighborhood of the predicted position of a given process in the reconstructed landscape and the local neighborhood of the same process in the reference landscape. The local neighborhood is defined by the 100-closest neighbors in both landscapes, and the overlap between predicted and known neighborhoods is measured by the Jaccard index. A perfect neighborhood overlap indicates a high match with an index value of 1. These evaluations were performed at increasing removal percentages (from 0.1 to 90) and removing gene sets randomly 20 times in each case. Average evaluation scores are shown in **Fig.S4c**.

### Neuronal specificity score

The neuronal specificity score was computed by multiplying the rank of the *lbSpec* score in the reference cell types and the rank of the average expression of genes in excitatory and inhibitory neurons. The score was then normalized by dividing it by the total number of genes squared.

### Expression breadth deviation

The expression breadth (B) deviation between two species was derived by computing the mean absolute B level difference for each L value across all genes.

## Data availability

Precomputed reference L-B lines, landscapes, and specificity measurements are included in the accompanying R package *GeneSLand*. References include data for human tissues, cell type, and cortical neuronal subtypes; and data for cortical subtypes of adult humans, chimpanzees, rhesus macaques, and common marmosets. Corresponding expression profiles are reported in Supplementary data.

## Code availability

GeneSLand implementation in R is available at https://github.com/davilavelderrainlab/GeneSLand.

## Acknowledgements

We thank members of the davila-velderrain group for useful discussion. This research was supported by the Human Technopole Foundation supporting J.D.-V.

## Author contributions

E.B. performed all computational analysis and implemented the R package. J.D.-V. conceived and directed the study. All authors designed, organized, and wrote the manuscript.

